# Declining RNA integrity in control autopsy brain tissue is robustly and asymmetrically associated with selective neuronal mRNA signal loss

**DOI:** 10.1101/2021.09.07.459326

**Authors:** Eleanor S. Johnson, Kendra E. Stenzel, Sangderk Lee, Eric M. Blalock

**Affiliations:** Department of Pharmacology and Nutritional Sciences, University of Kentucky, Lexington, Kentucky, United States of America

## Abstract

RNA integrity numbers (RINs) are a standardized method for semi-quantification of RNA degradation, and are used in quality control prior to transcriptional profiling analysis. Recent work has demonstrated that RINs are associated with downstream transcriptional profiling, and correction procedures are typically employed in bioinformatic analysis pipelines to attempt to control for RIN’s influence on gene expression. However, relatively little work has been done to determine whether RIN’s influence is random, or is specifically targeted to a subset of mRNAs. We tested the hypothesis that RIN would be associated with a robust transcriptional profile seen across multiple studies.

To test this, we downloaded subsets of raw transcriptional data from six published studies. We only included control, non-pathological post-mortem human brain tissue (n = 383 samples) in which independent subjects’ RIN values were also reported. A robust set of mRNAs consistently and significantly correlated with RIN across multiple studies, appearing to be selectively degraded as RIN declines. Many of the affected gene expression pathways are related to neurons (e.g., vesicle, mRNA transport, synapse, and mitochondria), suggesting that neuronal synaptic mRNA may be particularly vulnerable to degradation. Subsequent analysis of the relationship between RIN and vulnerable mRNA expression revealed most of the decay occurred over a relatively narrow RIN range of 7.2-8.6, with RIN values > 8.6 showing a ceiling effect, and those < 7.2 showing a floor effect on gene expression. Our data suggests that the RIN effect is pathway selective and non-linear, which may be an important consideration for current bioinformatic RIN correcting procedures, particularly in datasets in which declining RIN is confounded with a pathology under study (e.g., in Alzheimer’s disease).

## Introduction

Transcriptional profiling in human brain tissue reveals that the gene signatures of aging and neurodegenerative diseases are robust and consistent across different independent samples using measurement platforms across different labs (1, 2). However, disease-states and aging are not the only factors possibly impacting gene expression in transcriptional profiling. Tissue treatment, post-mortem interval (PMI), pH, etc. also have an impact on gene expression (3-6). Often, these factors are assumed to impact gene expression, since they damage RNA quality.

RNA quality, as a measured through RIN, is known to impact transcriptional profiling (7-10). While there are many ways to determine RNA quality (i.e. gel optical density, NanoDrop, denaturing agarose gelelectrophoresis), but RNA integrity numbers (RINs) have become a standard in the last fifteen years (11-13). This technology relies on several measures including the total RNA ratio (fraction of the area in the 18S and 28S region compared to the total area under the curve), 28S peak height, fast area ratio (fast area compared to the total area), and the marker height (11). In addition, the program used to determine RINs also provides the RNA area under the curve, RNA concentration, and rRNA ratio (28S/18S). It should be noted that though mRNA is typically the focus of most transcriptional profiling studies, RIN heavily relies on rRNA to infer the quality (11, 13). Despite its prevalence and the importance of RNA integrity for transcriptional profiling, RINs are not often reported for individual subjects within published data. This is problematic, since often the tissue collection, storage, and handling can influence RNA quality (6, 10). Often, human tissue samples are more variable and frequently have lower RINs than those in experimental animal studies (See Results).

Since RNA quality may impact gene expression, particularly in humans, multiple groups have published bioinformatic tools to attempt to correct for the influence of RNA degradation on gene expression. Three approaches for correcting include Surrogate Variable Analysis (SVA) (14), quality SVA (qSVA) (9), and regression (10, 15, 16). SVAs use the gene expression data to determine which genes are impacted by sources of variability, depending on the metavariable being corrected. The factors are then used in a linear model to adjust for noise (14). However, Jaffe et al. was concerned that this approach might include false-positives. Therefore, they created their own model, qSVA, which uses results from a defined independent tissue degradation experiment to identify genes most affected by degradation and correlate them in the experimental dataset based on factors calculated in the independent dataset (9). Regression approaches are most commonly used to correct for RIN (10, 15, 16) since RNA degradation can influence genes and pathways at different rates (9, 10).

However, these approaches may be problematic for two reasons. First, if RIN values above a certain threshold are considered ‘safe’ (i.e., do not influence gene expression), then defining that safe threshold would be important, as the simplest approach would be to retain samples that exceeded the safety threshold. Further, attempting to control for a metavariable that is not associated with signal can cause artificially inflated variance (17), thereby distorting the processed signal. Second, as Gallego-Romero et al. reported, half of the differentially expressed genes after 84 hours of room temperature degradation appeared to show increased expression after RIN correction, but this increase was a distortion caused by the RIN correction procedure itself (10). Thus, establishing a safe threshold for RIN, one above which transcriptional profiling data can be processed without correcting for RIN could be of use. Further, it is important to appreciate whether the influence of RIN 1) is randomly distributed across the transcriptome, and therefore would not replicate in samples in a study or across studies from different labs; or 2) is focused on certain genes and pathways. Finally, whether the RIN effect is exerted across the full spectrum of RIN values or is constrained to a narrow range would be important to know as regression tools are more well-suited to the former than the latter case.

If a condition such as neurodegenerative disease is associated with lower RINs, then the two variables would be confounded, complicating attempts to control for one variable without influencing the other. Indeed, this is a common issue, as prior work (9, 15, 18) has shown that various insults are associated with significantly lower RINs in brain tissue. For instance, one study investigating the influence of pre- and post-mortem variables on RIN (18) found that ante-mortem variables such as agonal state, coma, and artificial ventilation had a cumulative downward influence on RIN, as did a diagnosis of Alzheimer’s disease (no RIN corrections were employed, likely because of the presence of this confound). Another study determined that RIN scores decreased with a diagnosis of schizophrenia (9). Noting that multiple-regression RIN correction approaches would be confounded, the authors applied their novel RIN correction procedure, qSVA, that estimated decay based on observations in independent degradation datasets. Yet here, RIN-associated mRNA declines in the schizophrenic brain exceeded the correction, and therefore exceeded the amount of decline seen in control tissue (9). It is also important to note that the schizophrenic brains did not have a longer PMI than their controlled counterparts, yet had lower RINs, suggesting that some process other than PMI, and possibly influencing different pathways, plays a role. This indicated not only that schizophrenia was associated with a decline in RIN, but that the mRNA species associated with RIN in schizophrenia were targeted more aggressively than in control tissue. Finally, the last study established that there is a significant decrease in RIN with Alzheimer’s disease (AD) (15). They also identified that a standard regression-based RIN correction procedure removed the well-established AD-effect from the transcriptional profile. While Jaffe et al., postulate an interaction between RNA quality and cellular composition (9), Miller et al. concluded the difference in RNA quality might be due to poor handling of AD tissue (15).

Due to the assumption that most post-mortem factors impact RINs, problems with current RNA correction methods, and the potential of biological factors playing a role in RNA degradation, we examined the role RINs play in gene expression in microarrays. In the current work, we investigate if RINs in control, post-mortem human frontal lobe tissue impacts gene expression in a non-linear fashion, targeting specific genes and pathways, and the thresholds of this impact. In addition, we investigated the relationship of other metavariables with RIN and the effect of RIN on AD tissue to determine if they impact similar genes.

## Methods

### Identifying datasets and preparing cel files

The .cel files for the control subjects matching our criteria (see results) of the datasets were downloaded. Different datasets may have used different probe level algorithms in their published work, and many probe level algorithms are influenced by the total number of arrays selected for calculation (19). Further, in our approach, the total number of samples that would be used likely would change because our unique selection criteria would assess only control tissue from specified brain regions. These files were analyzed in R using the Robust Multi-array (RMA) (20, 21) function in the oligo package (22) from Bioconductor. Therefore, we ran the RMA probe level summarization algorithm independently on .cel files of each dataset. These estimates of gene expression were transferred to flat files in Excel, and Gene Symbols and Titles (based upon the updated GEO platform IDs under which the original datasets were published), as well as sample-specific metadata (e.g. RIN, Age, pH, PMI, sex) were annotated.

### Differences in RNA quality in brains across species

To determine if there was a significant difference in reported RNA quality between human, rat, and mouse brains as most assume (keeping in mind that because RIN values are used to triage subjects, there is likely some bias in these reports), studies were identified that matched the following criteria: 1) include control brain tissue (for humans, only frontal lobe); 2) have disambiguated RNA integrity numbers (RINs); and 3) use Affymetrix platforms. Once studies were found, the RIN scores were separated by species and an unequal Mann-Whitney U-Test quantitatively tested the hypothesis (observed anecdotally in prior work) that human autopsy samples show lower RIN score than those from experimental animals. Subsequent work focuses completely on human, post-mortem brain tissue.

### Pre-statistical processing and individual study analysis

Once the .cel files were analyzed, each study was independently processed as follows. We created signal intensity histograms to determine presence-call cutoffs (PCC). The PCC was determined by analyzing the signal intensity histogram to determine the ‘saddle point’ at which signal values transitioned from noise to reliable signals. Then, within each dataset, probe sets were retained for analysis if ≥ 50% of that probe set’s observations ≥ PCC. To retain unique gene symbol representations, if > 1 probe set was annotated to the same gene symbol, then the probe set with the highest average signal intensity was retained.

### Separate dataset approaches

#### Defining significant metavariable-correlated genes

Once the pre-statistically filtered, uniquely annotated, and reliably expressed probe sets were selected, Pearson’s correlation (r) tests between different metavariables (i.e. age, PMI, and pH), and between metavariables and expression signals were performed, and the associated p-values and false discovery rates (FDRs) calculated. A liberal p-value cutoff (α = 0.05) defined significant results to address the increased false negative rate anticipated to occur in subsequent cross dataset comparisons (23, 24). To investigate the potential influence of sex on RIN-associated gene expression levels, the males and females from the data-subset with the largest number of subjects underwent independent RIN-associated gene expression analysis. In this dataset, males still represented ∼3/4 of the total dataset. To address this statistical power imbalance, we also randomly resampled (1000 iterations) male subjects at the female n-level for comparison between sexes.

#### Comparing across separate datasets

The influence of metavariables on independent data-subsets was assessed with a post-hoc proportional change analysis followed by binomial testing to determine whether there was significant agreement. The significant common probe sets were run through the WEB-based Gene Set Analysis Toolkit (WebGestalt) (25) to statistically test functional categories in the Gene Ontology (26, 27) for overrepresentation (α = 0.05; GO Molecular Function, Cellular Component, Biological Process noRedundant; Redundancy reduction = affinity propagation).

### Unified dataset analysis

#### Approaches for unifying independent datasets/ removing batch effects

The ‘independent dataset’ approach described above treats each data set as an independent unit, therefore avoiding potential batch effects. However, the ‘independent dataset’ approach is also limited by the range of RIN values available in each study. To unify independent datasets, we tried three methods.

> Method 1) Within-dataset standardization. Within each dataset, each row’s data is standardized, then those standardized results are combined across datasets
>
> Method 2) Harmony. Using the Seurat package (28) within the Bioconductor (29, 30) subset of the R programming language, the Harmony function (31) was applied to use PCA-based metrics to subtract batch-based variance.
>
> Method 3) Mean-subtraction. Within each dataset, for each gene, the average gene expression for subjects with a RIN ≥ 8.3 were averaged. Then, that average was subtracted from all observations for that gene from that dataset.

#### Template analysis

Unified dataset gene expression data were placed into a single worksheet and correlated with 356 user-defined templates. These templates were used to fit the gene expression signal across RIN values. Correlation was performed for each gene with all 356 templates, and the gene was assigned to the template with which the gene correlated most strongly and exceeded a stringent correlation criterion (r ≥ |0.69|; p = 9.61 E-5). Then, the number of genes observed per template was quantified. To determine if the number of genes assigned to each template was greater than expected by chance, a resampling procedure (1,000 iterations) was performed to estimate the number of genes per template. The probability that the number observed in the actual data was significantly higher than number estimated in the resampled data was tested for significance using the Z-score. Pathway analysis on selective templates with significant genes proceeded as described for ‘separate dataset approaches.’

### Alzheimer’s disease gene relationship

In order to determine if pathways associated with Alzheimer’s disease (AD) are overrepresented among RNA degradation sensitive genes, two Alzheimer’s disease transcriptional profile studies were used (1, 15) (one examines the same tissue region as the present work (15), the second represents consensus AD signature across multiple brain regions from several labs (1)). From Miller et al., 2017 (15), AD-significant gene expression levels were identified based analysis of the original publication (p ≤ 0.05).

Hargis & Blalock (2017; (1)), a secondary data analysis study that summarized log2 fold-changes (L2FCs) from 5 AD brain transcriptional studies across 8 different brain regions was used. In the present work, a consensus list of AD-significant genes from that secondary data analysis was determined by calculating the average L2FC (± 99% confidence interval) across all 5 of those studies. Results were considered significant if the 99% confidence interval did not cross the ‘0’ L2FC line (e.g, 99% confidence the observation was upregulated or downregulated with AD). The binomial test was used to determine if the transcriptional profile of RIN-sensitive genes bore similarity to either AD profile (Miller or Hargis). The log_2_ fold changes (L2FC) for the difference between AD and control subjects for each gene in Miller et al. was then compared to the correlation between gene expression and RIN from study with the most the subjects. The average L2FC for each unique probe set from Hargis & Blalock was compared with the genes significant in all data-subsets used in the cross-data comparison.

## Results

### Overview of datasets

Using the Gene Expression Omnibus, we identified 6 datasets which met our criteria (disambiguated RNA integrity number [RIN] values; human frontal lobe tissue; and Affymetrix platform technology): GSE22521 (32), GSE25219 (33), GSE45878 (34), GSE46706 (35), GSE53987 (36), and GSE71620 (37) (Table 1).

**Table 1.**
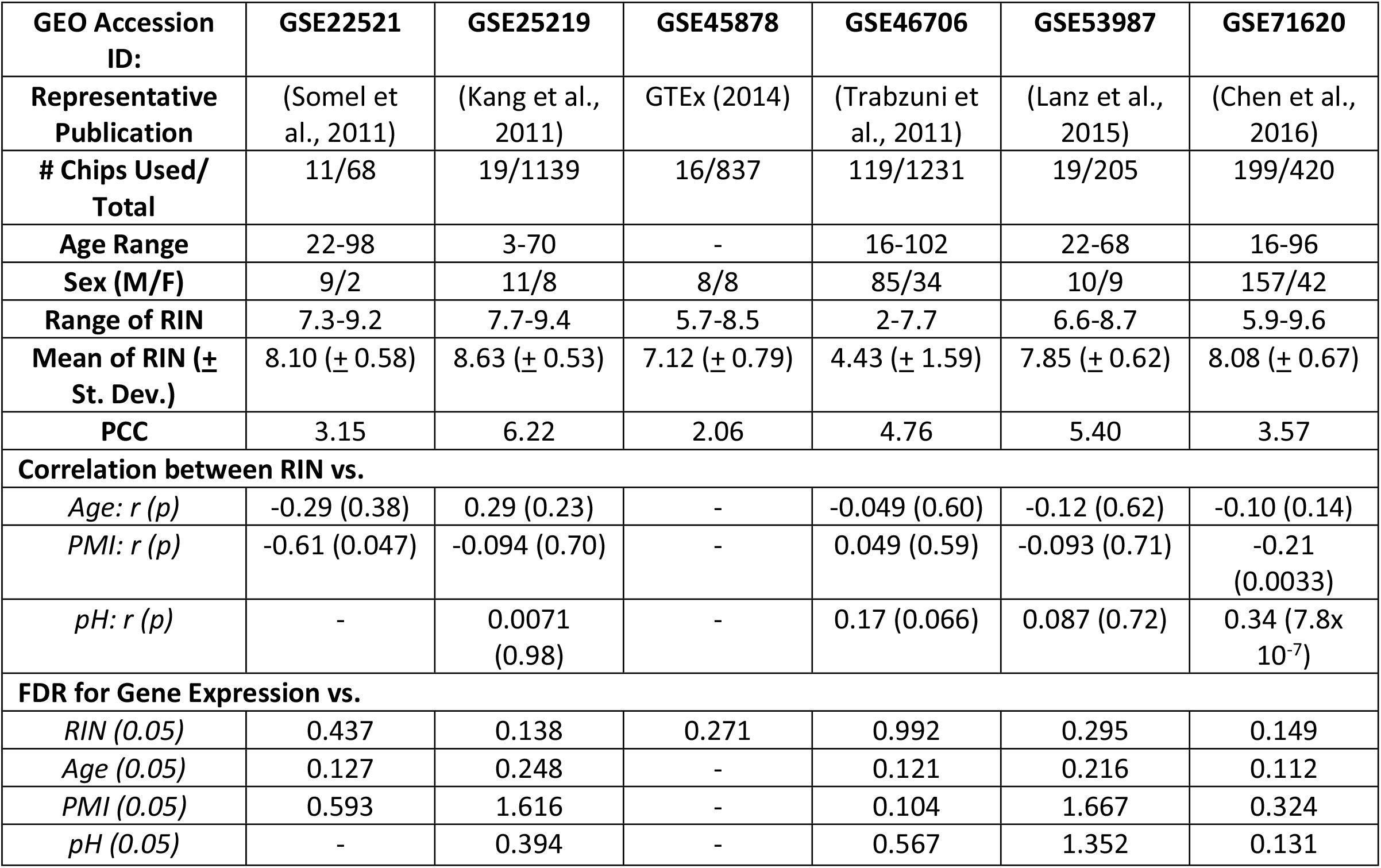
Data-subset information. *GEO Accession ID-* The Gene Expression Omnibus (GEO) identifier for each dataset; *Representative Publication-* peer-reviewed citation for each dataset; *#Chips Used/ Total-* The subset of the total chips within original dataset that met criteria for inclusion in the present work (e.g., from control subject frontal cortex-subsequent data in table is for subjects that met criteria); *Age Range-* age (in years); *Age and RIN:* Results of correlation analysis (Pearson’s test r and p-values) between age and RNA-Integrity Number; *Sex:* number of male (M) and Female (F) subjects; *Range of RIN, Mean of RIN* are as described; *FDRs for RIN, Age, PMI and pH-* False Discovery Rate (FDR) for the correlation between these metavariables and global gene expression.

| GEO Accession ID: | GSE22521 | GSE25219 | GSE45878 | GSE46706 | GSE53987 | GSE71620 |
| --- | --- | --- | --- | --- | --- | --- |
| Representative Publication | (Somel et al., 2011) | (Kang et al., 2011) | GTEX (2014) | (Trabzuni et al., 2011) | (Lanz et al., 2015) | (Chen et al., 2016) |
| # Chips Used/ Total | 11/68 | 19/1139 | 16/837 | 119/1231 | 19/205 | 199/420 |
| Age Range | 22-98 | 3-70 | - | 16-102 | 22-68 | 16-96 |
| Sex (M/F) | 9/2 | 11/8 | 8/8 | 85/34 | 10/9 | 157/42 |
| Range of RIN | 7.3-9.2 | 7.7-9.4 | 5.7-8.5 | 2-7.7 | 6.6-8.7 | 5.9-9.6 |
| Mean of RIN (± St. Dev.) | 8.10 (± 0.58) | 8.63 (± 0.53) | 7.12 (± 0.79) | 4.43 (± 1.59) | 7.85 (± 0.62) | 8.08 (± 0.67) |
| PCC | 3.15 | 6.22 | 2.06 | 4.76 | 5.40 | 3.57 |
| **Correlation between RIN vs.** |  |  |  |  |  |  |
| Age: *r* (*p*) | -0.29 (0.38) | 0.29 (0.23) | - | -0.049 (0.60) | -0.12 (0.62) | -0.10 (0.14) |
| PMI: *r* (*p*) | -0.61 (0.047) | -0.094 (0.70) | - | 0.049 (0.59) | -0.093 (0.71) | -0.21 (0.0033) |
| pH: *r* (*p*) | - | 0.0071 (0.98) | - | 0.17 (0.066) | 0.087 (0.72) | 0.34 (7.8x 10<sup>-7</sup>) |
| **FDR for Gene Expression vs.** |  |  |  |  |  |  |
| RIN (0.05) | 0.437 | 0.138 | 0.271 | 0.992 | 0.295 | 0.149 |
| Age (0.05) | 0.127 | 0.248 | - | 0.121 | 0.216 | 0.112 |
| PMI (0.05) | 0.593 | 1.616 | - | 0.104 | 1.667 | 0.324 |
| pH (0.05) | - | 0.394 | - | 0.567 | 1.352 | 0.131 |

From within each dataset, the .cel files that qualified for analysis (the qualifying “data-subset”- [tissue must be human frontal lobe tissue, must include disambiguated RNA integrity numbers (RINs), and must use Affymetrix platform technology) were downloaded and reanalyzed at the probe level using Robust Multi-array (RMA; (20, 21)) (Supp. Data 6-11). For each data-subset, a signal frequency histogram was plotted to remove low-intensity signal (Figure 1). Further, the gene annotations for each data-subset were downloaded from the platforms most recently associated with the studies on Gene Expression Omnibus (GEO) (38). GPL6244 (last updated March 5, 2020) annotated GSE22521; GPL5175 (last updated February 18, 2019) annotated GSE25219 and GSE46706; GPL16977 (last updated September 2, 2014) annotated GSE45878; GPL570 (version updated June 16, 2016) annotated GSE53987; and GLP11532 (last updated November 8, 2016) annotated GSE71620. For each data-subset, if more than one row of data was annotated to the same gene symbol, then the row with the highest mean signal intensity was retained for further analysis.

**Figure 1.**
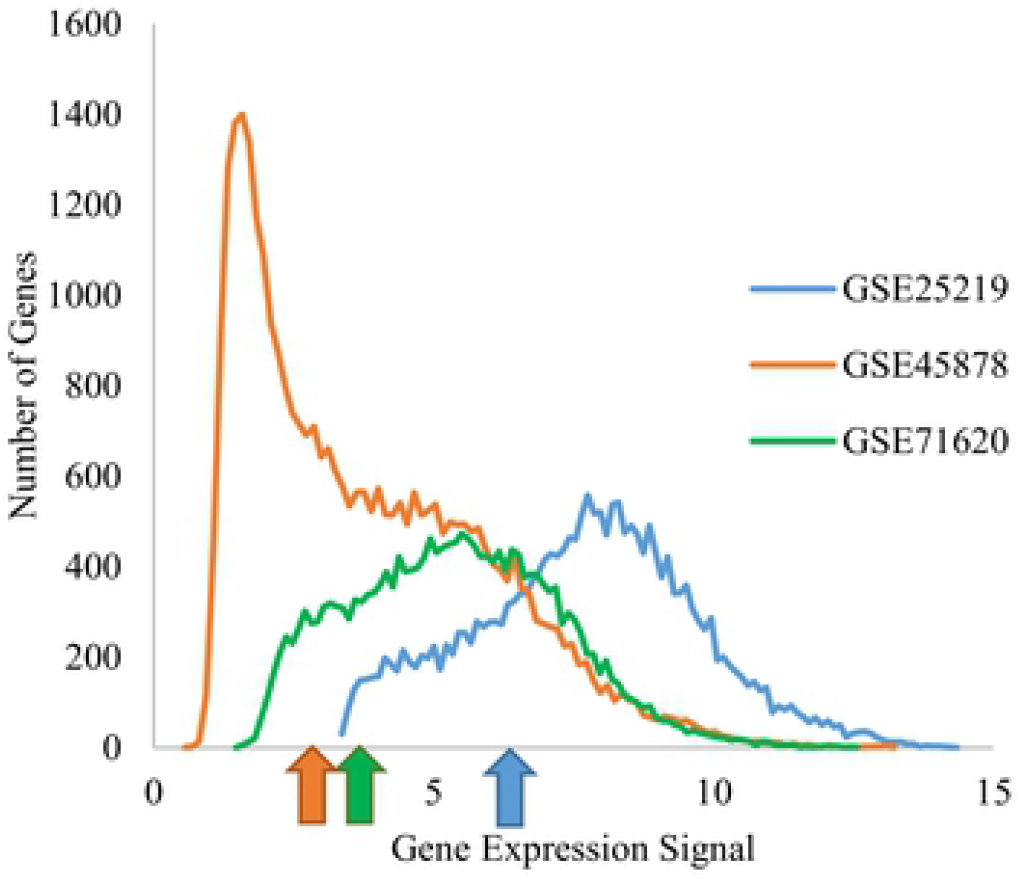
Presence cut-offs established in a dataset-dependent fashion. Histograms of the average gene expression for datasets determine the cut-off value used. The gene expression signal varied significantly between the datasets. This prevented us from a single standardized cutoff value across all studies and combining the datasets, so each study had an independent cut-off value. Arrows represent the cut-off value for study of the same color.

For this secondary data analysis, two approaches are feasible. The first, and more conservative, approach would be to analyze each data-subset independently for correlation to RIN, and then, to compile the statically significant results from these independent tests. Using this approach, agreement among multiple independently analyzed datasets regarding RIN-sensitive genes could be used to test for effects. However, its conservative nature is both a strength and a weakness, reducing the likelihood of statistical false positives at the expense of increasing the likelihood of statistical false negatives. A second, more liberal approach would be to combine the data from these disparate sources to create a single dataset. This approach would have the advantage of increased discovery power but would also be more vulnerable to false positives, and would only be feasible in the absence of batch effects. To determine whether the first (‘independent dataset’) or secondary (‘merged dataset’) approach is more feasible for the initial analysis, a principle component analysis (PCA) was constructed. The PCA shows an consistent effects. This approach identifies robust effects and is less vulnerable to dataset-based batch intense batch effect among the different datasets (Figure 2A), strongly suggesting the ‘independent dataset’ approach would be more feasible for the data prior to any batch removal procedures.

**Figure 2.**
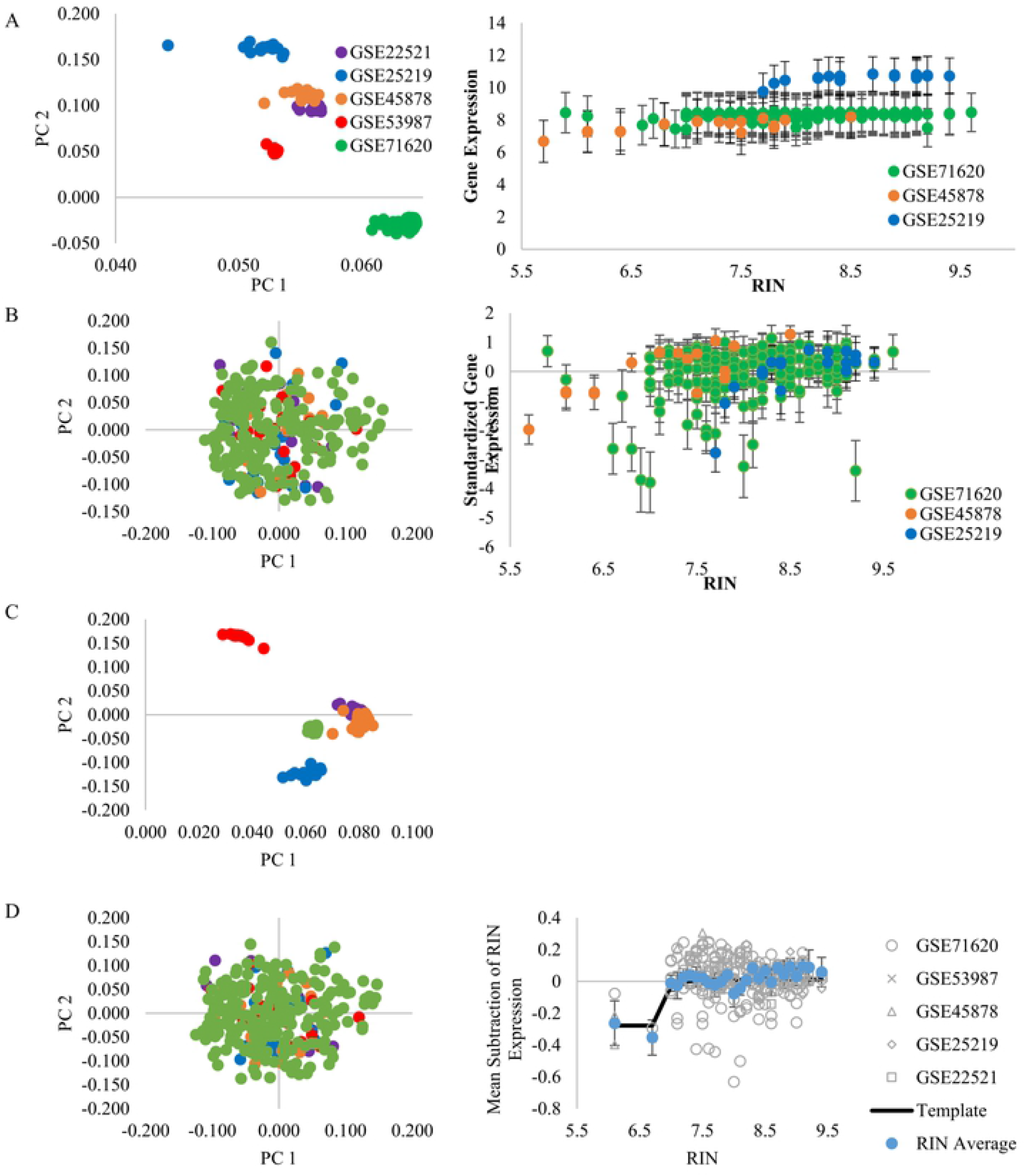
PCA plots and monovalent inorganic cation transmembrane transport pathway expression. The top 2,000 variable genes were used in the principle component analysis (PCA). The different datasets are distinct and separate from each other, not allowing for a single analysis. Genes for the monovalent cation transmembrane transport pathway (Table 4) were used to determine how gene expression decreased as RNA degrades. Here, we show the RMA values (A); a standardized gene expression (B); Harmony (C); and the mean-subtraction (D). Therefore, the mean subtraction approach (D) appeared to both remove the batch effect, while preserving original study deflecting points in the merged data. The raw data and Harmony did not remove the batch effect, and therefore did not allow for a merged data across datasets. The standardization method had each unique data-subset beginning to decline at different points, which may be due to the centering each dataset at zero, regardless of RIN range.

### Differences in RNA quality in brains across species

While reading prior studies, it came to our attention that no one had supported the claim that animal RINs were typically higher than those of humans. Therefore, we found ten studies which met our criteria (control brain tissue; disambiguated RINs; and Affymetrix platform technology) (32-37, 39-42) (Table 2).

**Table 2.**
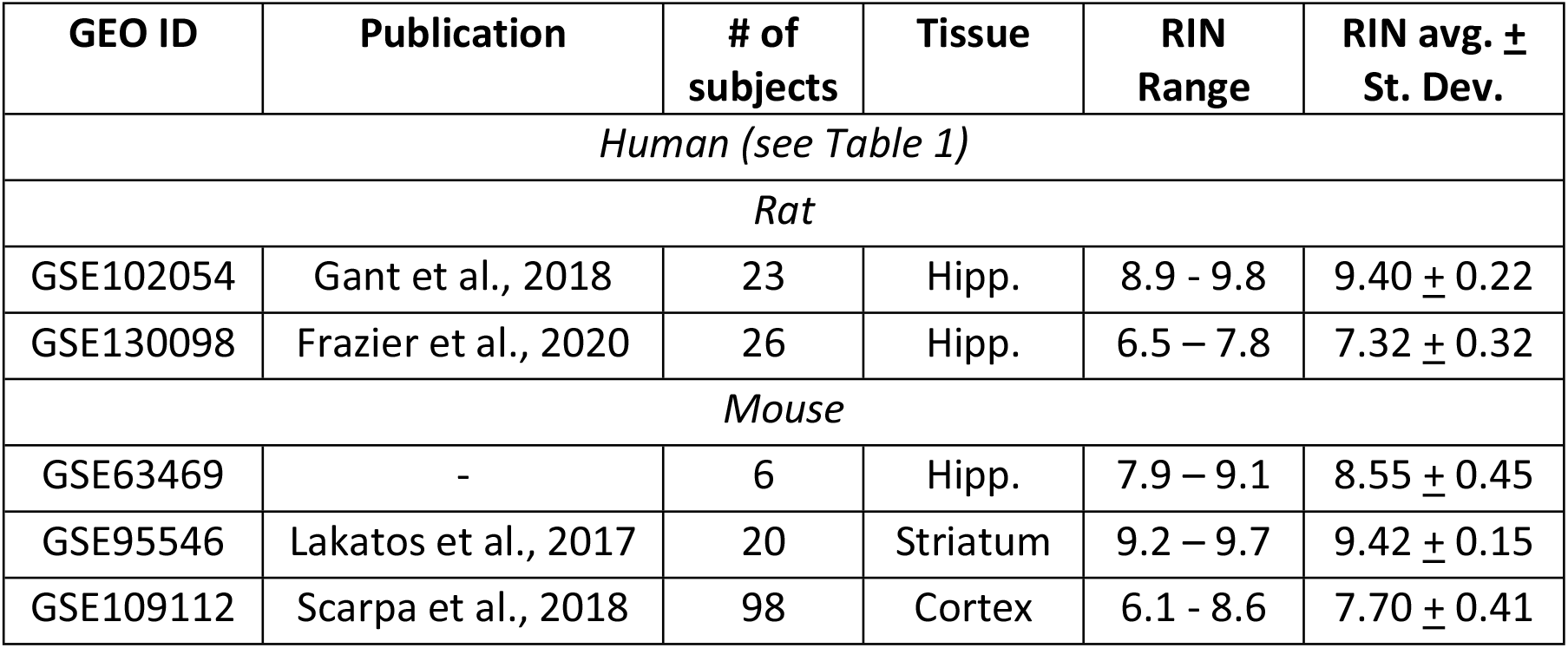
RIN comparisons across species. GEO Accession ID-The Gene Expression Omnibus (GEO) identifier for each dataset; Representative Publication-peer-reviewed citation for each dataset; # of subjects-The number of subjects from a subsection that met our criteria in the present work (e.g. from human control subject frontal cortex-subsequent data in table is for subjects that met criteria) or a single tissue type from each dataset (rodents); Tissue-Reported tissue type analyzed; RIN Range, RIN Avg. + St. Dev. as described. Hipp.-hippocampus.

| GEO ID | Publication | # of subjects | Tissue | RIN Range | RIN avg. ± St. Dev. |
| --- | --- | --- | --- | --- | --- |
| *Human (see Table 1)* |  |  |  |  |  |
| *Rat* |  |  |  |  |  |
| GSE102054 | Gant et al., 2018 | 23 | Hipp. | 8.9 - 9.8 | $9.40 \pm 0.22$ |
| GSE130098 | Frazier et al., 2020 | 26 | Hipp. | 6.5 – 7.8 | $7.32 \pm 0.32$ |
| *Mouse* |  |  |  |  |  |
| GSE63469 | - | 6 | Hipp. | 7.9 – 9.1 | $8.55 \pm 0.45$ |
| GSE95546 | Lakatos et al., 2017 | 20 | Striatum | 9.2 – 9.7 | $9.42 \pm 0.15$ |
| GSE109112 | Scarpa et al., 2018 | 98 | Cortex | 6.1 - 8.6 | $7.70 \pm 0.41$ |

When disambiguated RINs are grouped by species (human, rat, and mouse), there is a significant difference between human and rat (unequal Mann-Whitney U-Test p = 6.10E-4) and human and mouse (unequal Mann-Whitney U-Test p = 1.63E-6), but not between rat and mouse (unequal Mann-Whitney U-Test p = 0.301).

### Metadata correlation analysis of individual datasets

Pearson’ correlations between published gene expression values and metadata (RIN, pH, post-mortem intervals [PMI], age) were calculated for each individual dataset where available (Table 1; Figure 3). 0/ 5 datasets had a significant (p ≤ 0.05) age-to-RIN correlation (Figure 3A), while 2/ 5 had significant negative PMI-to-RIN correlations (Figure 3B). 1/ 4 datasets had a significant positive pH-to-RIN correlation (Figure 3C). Surprisingly, this lack of a robust relationship between RIN and PMI, or RIN and metadata traditionally associated with tissue quality suggests these measures may serve as poor proxies for estimating RNA quality.

**Figure 3.**
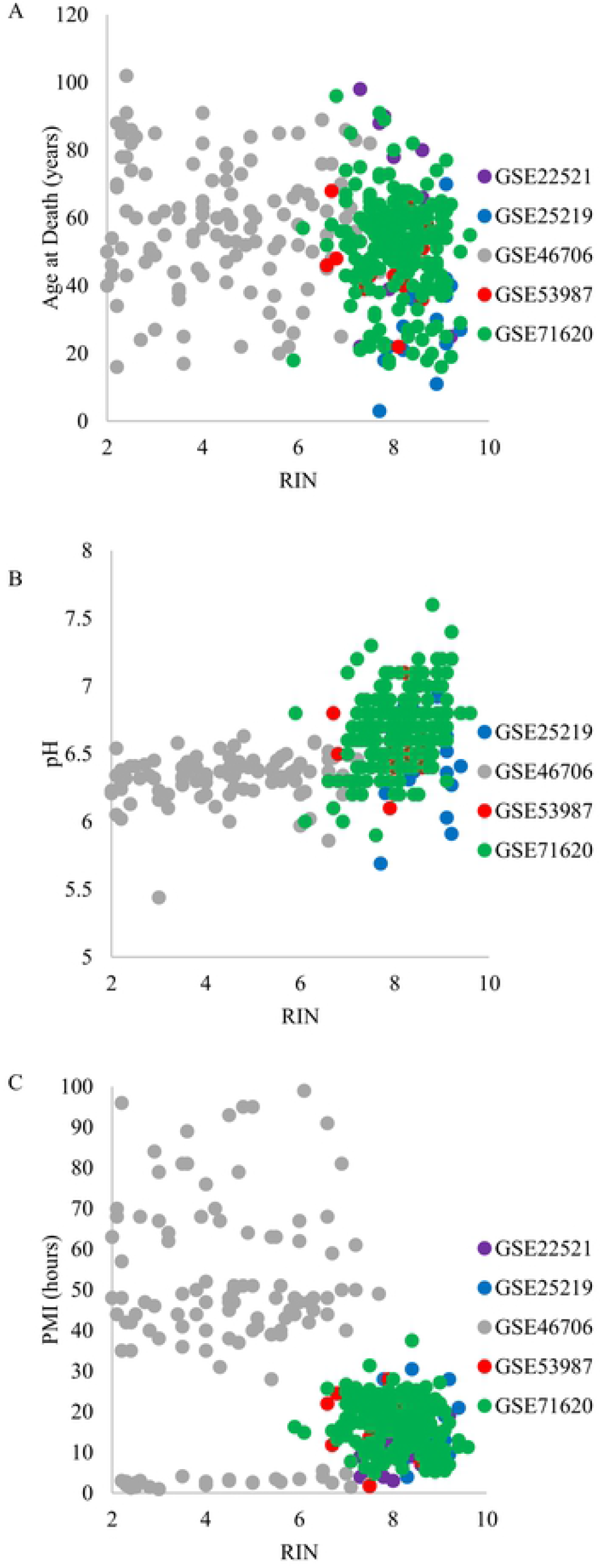
Metadata correlations to RIN. To determine if RNA integrity numbers (RINs) were significant correlated to any of the other commonly described metavariables, RIN was correlated to Age (in years) (A); Post Mortem Interval (in hours; PMI) (B); pH correlated to RIN (C).

However, just because metadata do not correlate with RIN scores does not mean they do not explain some degree of gene variability in their own right. To test this, the false discovery rates (FDRs) were calculated for the correlation between gene signal and: RINs; pH; PMI; or age (Table 1). Aging showed a strong and consistent influence on gene expression (average FDR = 0.165; 95% Confidence Interval [CI] = [0.110-0.220]) (32, 43-45). RIN appeared to have the second most prominent influence (average FDR = 0.380; 95% CI = [0.125-0.636]), while the influence of pH (average FDR = 0.611; 95% CI = [0.0960-1.13]) and PMI (average FDR = 0.861; 95% CI = [0.218-1.50]) were less stable. It should be noted that there are constraints on the data, determined by the original studies, which may be influencing the FDRs for RIN, pH, and PMI. Because our study focuses on the effects of RNA degradation, we focused on data-subsets with the lowest FDRs (indicating statically effects that were less likely to be false positives) for RIN (GSE25219, GSE45878, and GSE71620) for further analysis.

Using the dataset with the most subjects, GSE71620, we investigated potential sex difference in RIN sensitivity. When separated into males and females, males had 3650 RIN-correlated genes while females had 3251 genes. However, there was an imbalance between the numbers of male and female subjects (157 males/ 42 females; male RIN = 8.08 ± 0.630; female RIN = 8.09 ± 0.808), which likely influenced statistical discovery power. To estimate the potential influence of the differing number of male and female subjects, a random resampling approach was used. The 157 male subjects were randomized (1000 iterations) to determine, on average, what number of RIN-sensitive genes would be identified if the male subjects only had 42 subjects (the same as the female subset). The average and standard deviation of the resampling iterations was plotted along with the observed value of significant genes in females. When comparing males and females at the equally powered ‘42 subject’ level, the females discovered a non-significant (but trending) greater number of genes (3251 compared to 1719 [± 1065] resampled males, Z-test p = 0.075; Figure 4A).

**Figure 4.**
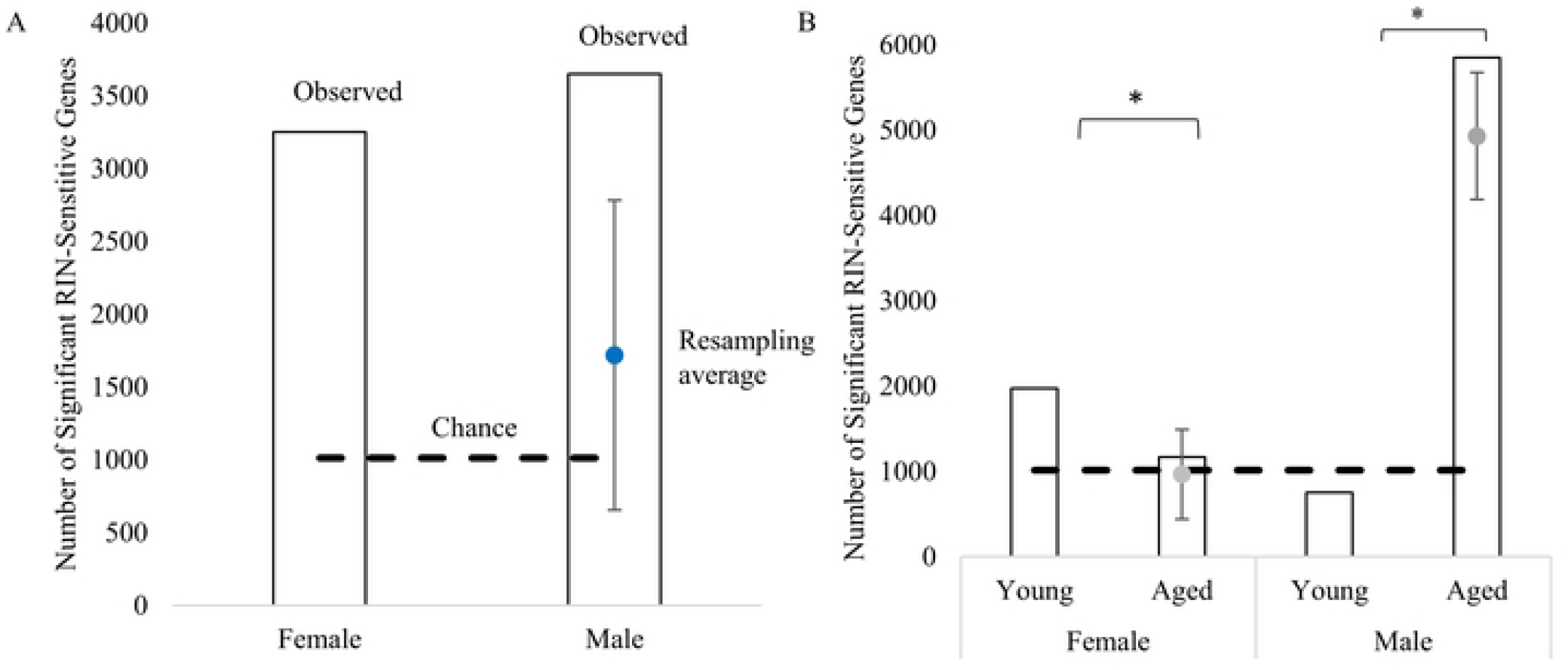
Sex effect on RNA degradation. To equalize the estimated power between the number of male and female subjects in GSE71620, and young and aged for each sex, the group with the higher number of subjects was randomly resampled at the same n as the group with the lower number of samples. The number of significant genes for males and females (A). The blue dot represents average number of significant genes from 1000 resampling iterations. The hashed black line indicates the number of genes expected by chance (1013; A). The number of significant genes for young and aged males and females (B). The hashed black line indicates the number of genes expected by chance (1013). The gray dots represent the average number of significant genes found when the aged male and female groups were resampled at the same power as the young groups (n = 66, n = 14 respectively; B). * < 0.05.

The females were also separated into young (n = 14; 16-46 years old; RIN = 8.32 ± 0.714) and aged (n = 28; 50-96 years old; RIN = 7.96 ± 0.840) groups and the aged subjects were randomized in the same manner. The young females had a significantly greater number of RIN-sensitive genes compared to their older counterparts (1972 young compared to 966 [± 526] resampled aged females, Z-test p = 0.028; Figure. 4B). The males were separated into young (n = 66; 17-49 years old; RIN = 8.09 ± 0.651) and aged (n = 91; 50-89 years old; RIN = 8.08 ± 0.617) groups and the aged group was randomly resampled, as well. The young males had a significantly lower number of RIN-sensitive genes compared to their older counterparts (755 compared to 4932 [± 755] resampled aged males, Z-test p < 1.0E -5; Figure. 4B).

Almost all of the significant genes in the aged male group had a positive correlation with RIN (total significant genes = 5853; positive significant genes = 4362; binomial test p < 1.0 E -15). To determine if this was by chance, the correlation was validated for these genes using in an additional study, GSE45878. GSE45878 contained 4356 genes out of the 4362 positive genes, out of which 2781 had a positive correlation between RIN and gene expression (binomial test p < 1.0 E -15). This indicates there is robust positive correlation between aged males’ gene expressions and RIN.

### Cross dataset analysis using individual datasets

A common list of total genes was determined from GSE25219, GSE45878, and GSE71620. This total gene list was comprised of probe sets annotated to unique gene symbols with sufficient signal intensity in all three studies (15,735 ‘total’ genes). Using a P-value cut-off for the correlation between the RIN and gene expression signal (p < 0.05), the number of genes significantly correlated with RIN in all 3 data-subsets was significant (609 compared to the expected 209; p < 1.0E-15; Binomial test; S1 Data) (Figure 5).

**Figure 5.**
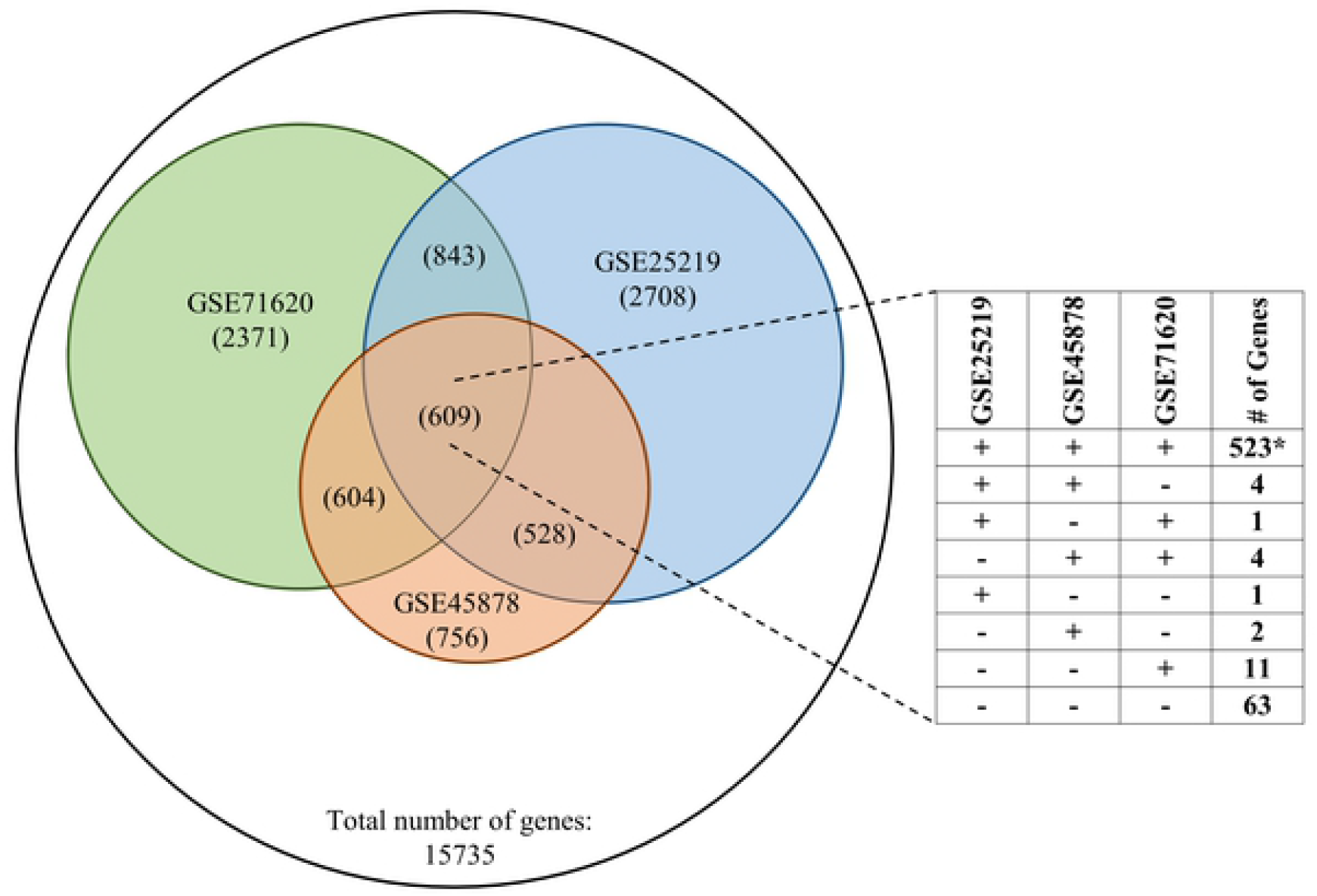
Overlapping significant genes between 3 datasets, and direction of significant genes. Using GSE25219, GSE45878, and GSE71620, a common list of 15,735 genes was determined. The number of significant genes in each study and the overlap was determine. The expected number of genes significant in all 3 datasets was expected to be 209. *Left 3 Columns*-the direction of the RIN-correlation for the study listed in the top row; positive correlation (+), negative correlation (-)). *# of Genes*-number of genes in each category. * p ≤ 0.05.

Further, this overlapping agreement showed a strong direction-of-correlation agreement. The overlapping genes significant in all three datasets could have fallen into one of eight potential categories based on the correlation direction between RIN and gene expression (Figure 5). The subset of genes positively correlated with RIN in all three data-subsets (523/ 609) was highly significant (p ≤ 1.0E-15), suggesting a consistent group of genes are robustly associated with RIN levels in multiple independent subjects. We validated the direction of change for these robust RIN-sensitive genes by comparing the correlative direction in a fourth dataset, GSE53987. Out of the 512 robust RIN-sensitive genes, 400 agreed in direction (binomial test: p < 1.0E-15; S2 Data) in GSE53987. Biological pathway overrepresentation analysis (WebGestalt) of these 523 robustly RIN-sensitive genes revealed consistent and selective pathways impacted, with the prominent positive correlation indicating a general decline in gene expression levels as RIN levels decreased (Table 3). These pathways appear to be related to vesicles, transporters, mitochondria, and synapses. This indicates mRNA housed in neurons may be particularly vulnerable to RNA degradation.

**Table 3.**
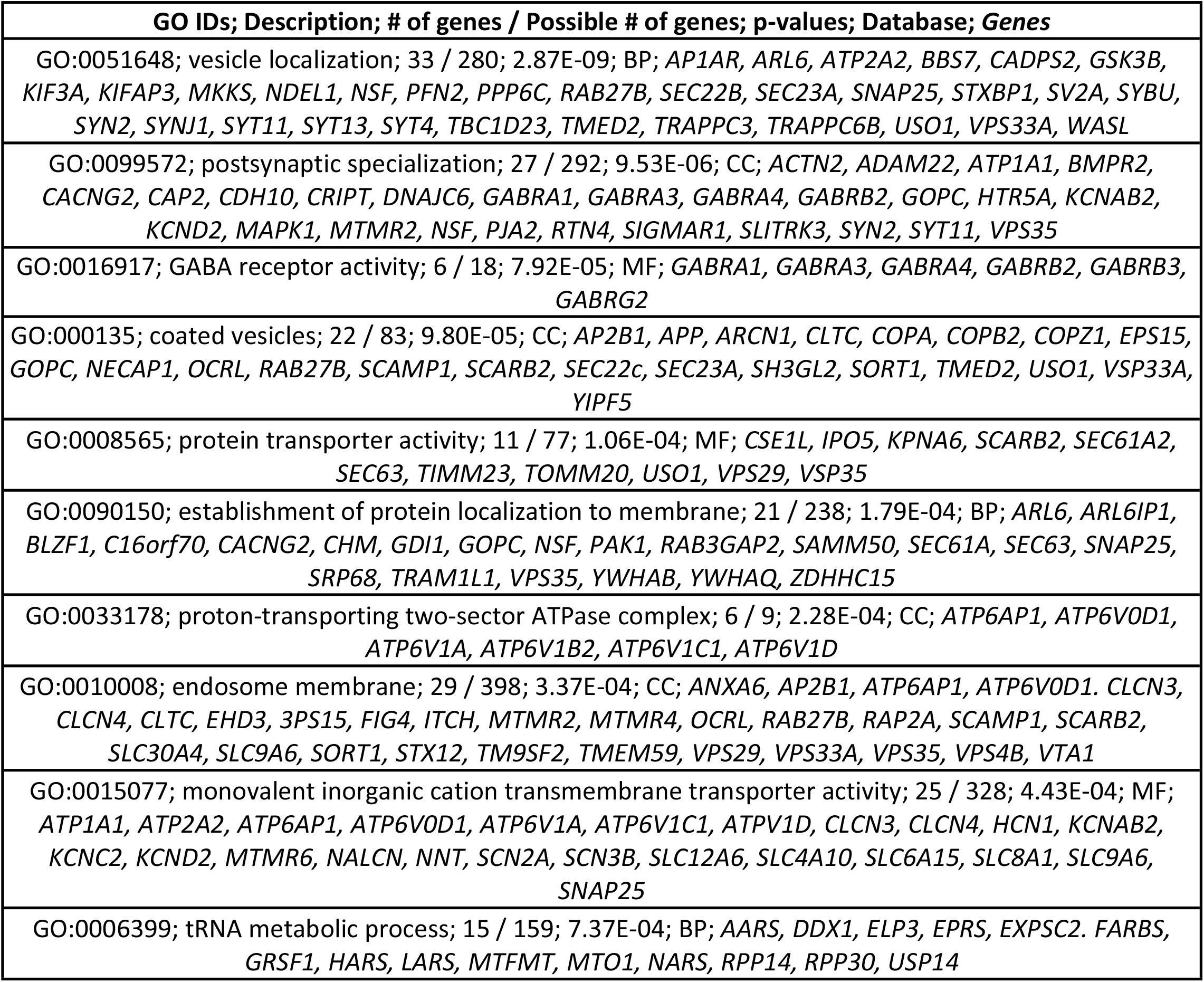
RIN-sensitive pathways. *Description-* name of the overrepresented pathway; *# of genes*-number of genes present in the group out of the 15,735 common list genes; *p-value*-p-value of finding this pathway by chance from our list; *Database*-functional group the pathway is associated with (Biological Process [BP], Cellular Component [CC], or Molecular Function [MF]); *genes (italized)*-list of RIN-sensitive gene symbols associated with pathway.

One consistent finding from the individual data-subsets analysis is the smaller r-values with larger data-subsets. When the data-subsets contain < 20 subjects, the maximum and minimum r-values (min r = - 0.8497; max r = 0.8737) are more extreme than the data-subsets with > 100 subjects tend to have a lower |r-values| (min r = - 0.3773; max r = 0.3343). While the higher-powered data-subsets do not need higher r-values to show significance, it indicates the Pearson correlation’s assumption of a linear relationship between gene expression level and RIN may not hold. This leads us to that the hypothesis the relationship between gene expression and RINs may not be linear.

### Merged datasets analysis

Using the independent dataset analysis approaches described above, we found a robust set of RIN-sensitive genes across multiple studies. However, this approach is limited by different ranges of RIN values in each dataset, and by the assumed linear relationship between RIN and gene expression that is implicit to the Pearson’s correlation test. To address these issues, we employed two strategies: merging datasets to get a more inclusive range of RIN values; and using a template correlation approach on this merged dataset to evaluate potentially more complex relationships between RIN and gene expression.

#### Approaches for merging multiple datasets

In order to merge the datasets, the batch effects (Figure 2) of each dataset were addressed. Three different approaches (described below) were considered.

1. Within-dataset standardization. Gene expression values from GSE25219, GSE45878, and GSE71620 were used. For each dataset, gene values were Z-scored for each subject. The PCA plot using this standardized approach shows the successful removal of the batch effect (Figure 2B). However, calculating the standardized average per dataset using 25 significant genes from one of the robustly overrepresented pathways, monovalent inorganic cation transmembrane transporter activity (Table 3), reveals an issue (Figure 2B). Each dataset shows the same significant positive correlation between gene expression and RIN, but different deflection points. GSE25219 begins its downward trend around a RIN of 8; GSE45878 begins to decrease around a RIN of 7; and GSE71620 begins to fall around 7.5 (Figure 2B). This shows that standardization effectively removes the batch effect, but centering at 0 biases the deflection range within the individual dataset’s RIN range in the combined dataset approach, so that it is not appropriate to compare across studies.
2. Harmony. As a second attempt to remove batch effects from multiple datasets, the Seurat package with the Harmony algorithm (31) in R, which uses multi-dimensional PCA-based variance to remove batch effects from disparate datasets, was used. However, this approach did not appear to substantially reduce the batch effect compared to that observed in the original datasets (Figure 2C).
3. Mean-Subtraction. As a third approach to remove the dataset-dependent batch effect while retaining a less-biased ‘RIN deflection point’ estimate, we used a signal subtraction strategy. It is well-understood hybridization-dependent mRNA quantification technology is strongly dependent on probe design (e.g., G-C content) (46, 47). Subtracting average signal intensity from each gene within each study would effectively address this probe-based variance, and it is this probe-based variance that likely contributes substantially to the batch effect. However, in the present work, signal values are influenced by RIN, and different datasets have different RIN ranges. Therefore, subtracting the row average for each gene from each dataset would be less effective than conditionally subtracting only the average of signal from samples for which the RIN exceeded some cutoff. That cutoff should represent a RIN value above which the RNA is of such high quality, that RIN’s association with gene expression is not detectable. To estimate what that RIN cutoff might be, data from GSE71620 was used.

Number significant genes is plotted as a function of RIN (Figure 6), and each dot represents a step from a low (starting at 6.1) RIN to high RIN (ending at 9.4). For each step, the genes found to be significantly correlated with RIN at the prior step are subtracted. Thus, each step reflects the degree to which an increasing RIN range contributes to RIN-sensitive gene discovery. We found the correlation between RINs and gene expression is relatively flat from 6.1 to 7.1, increases markedly from 7.4 to 8.6, and additional new RIN-sensitive genes are rarely discovered by adding additional subjects with RIN > 8.6, suggesting a RIN cutoff of 8.3 would be sufficient for calculating a signal average relation for RIN influence.

**Figure 6.**
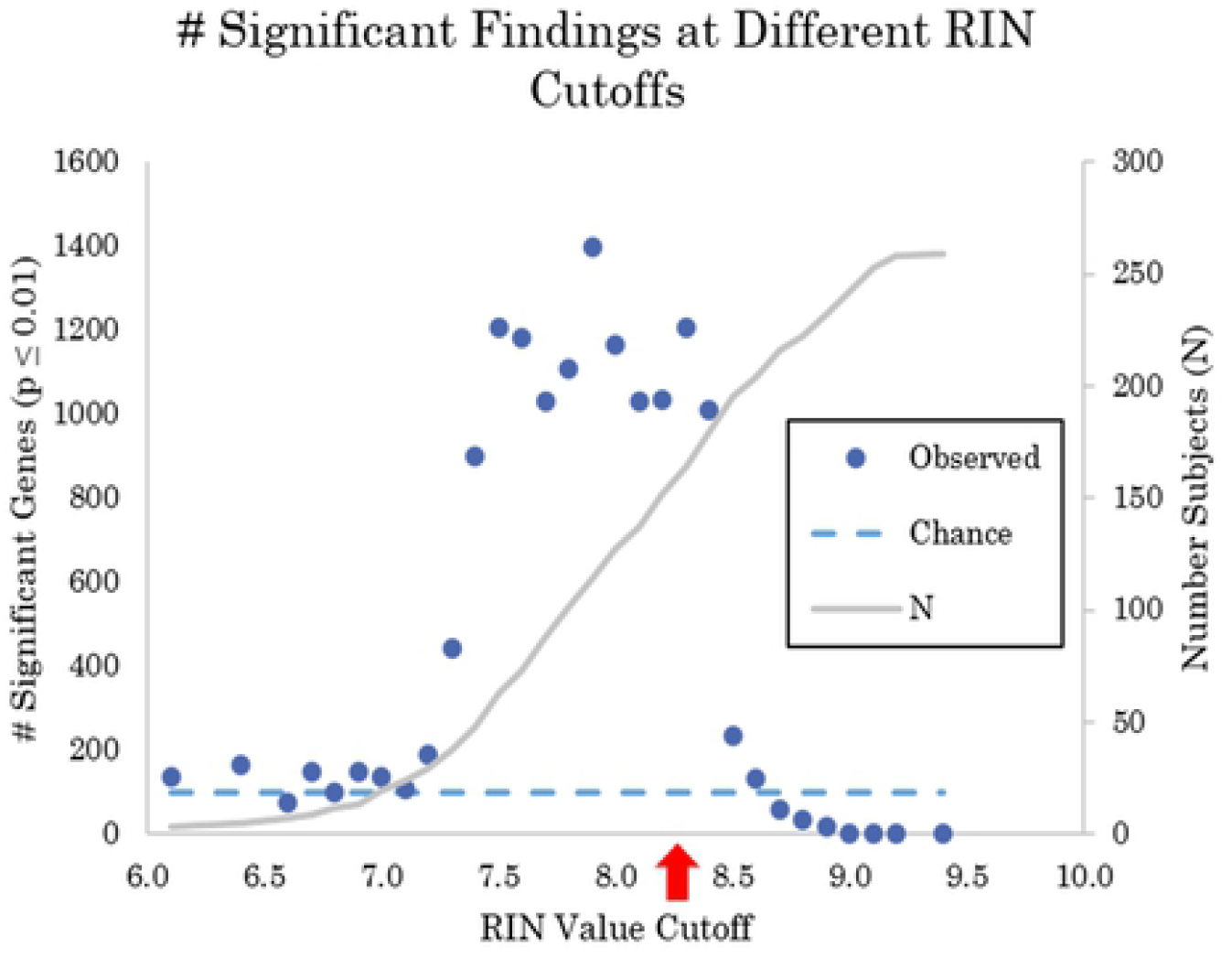
Number of significant genes associated with cumulative RIN. The number of genes whose expression levels were significantly correlated with RIN (Pearson’s test, p ≤ 0.01) are plotted as a function of RIN cutoff. For the first cutoff (RIN = 6.1) there were only 3 observations, and the results were similar to chance (dashed line). Between RIN values of 7 and 7.5, there is a marked increase in number of significant correlations. Similarly, between 8.3 and 8.5, there is a marked decrease in number of significant genes. This suggests 1) lower RIN values (6.1 to 7.1) are not appreciably correlated with gene expression (or the data is significantly underpowered), 2) there is a strong linear relationship between gene expression and RIN between 7.3 and 8.3, and 3) this relationship falls off sharply at RINs higher than 8.3 as gene expression plateaus. Taken together, these results suggest a sigmoidal relation relationship between gene expression and RIN, or an exponential decay obscured by decreased statistical power at the lowest RIN range.

Based on this, gene signal from subjects with a RIN > 8.3 appear less likely to contribute to the ‘RIN effect’ and their averages were calculated and then subtracted for each row of data in each dataset separately. Those ‘mean-subtracted’ datasets were then merged for subsequent analysis without a batch effect (Figure 2D).

### Template Analysis

Three hundred and fifty-six templates were designed that could potentially fit the gene expression signal across RIN values from the merged, mean-subtracted data (Figure 7; S3 Data). However, some of these templates were highly similar to one another. A Pearson’s correlation was run across the 356 templates, and those strongly correlated with one another were combined to simplify reporting. A single group of templates (from the S3 data, the linear climbs 6.1 to 7, 6.1 to 7.1, 6.1 to 7.2, exponential decline by 6.7, and plateau 6.1 to 6.7) were combined to make a “custom” template. Each gene was then correlated to all 352 templates, yielding 352 R values. Then, each gene was assigned to the template with which it most strongly correlated. If r ≥ |0.69| for its ‘best match’ template, then the gene was assigned to that template. 132/352 templates had genes assigned. Then, a resampling simulation (1000 iterations) was constructed to compare template assignment observed to estimated chance template assignment.

**Figure 7.**
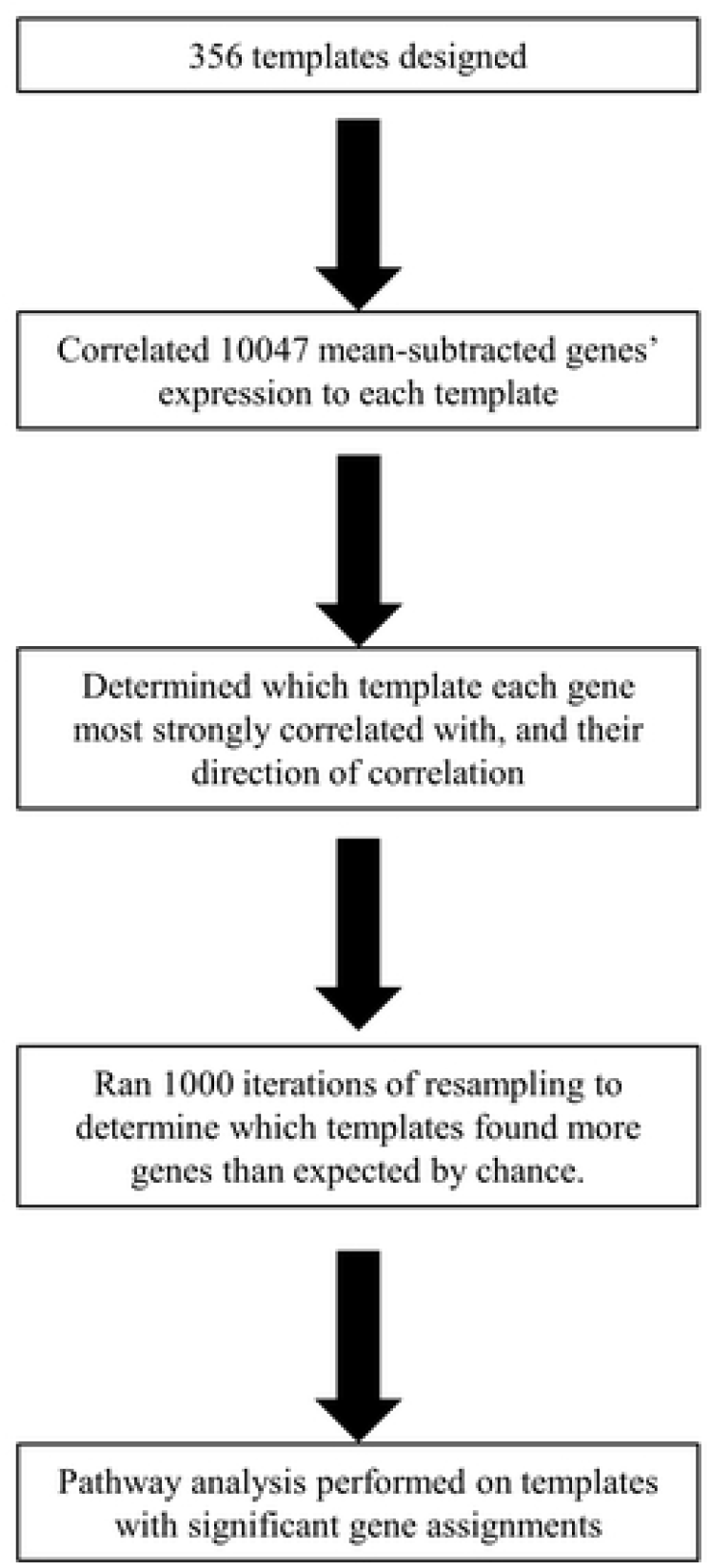
Flow chart of template analysis. Summarized the different steps taken during the template analysis.

Templates with significantly more gene assignments than estimated by chance (p < 0.05, Binomial test) were flagged for further analysis. Using this approach, 59 templates were identified as significant (Table 4, note that all template assigned genes are included in S1 Data) and the genes from the 6 templates with the largest number of gene assignments underwent pathway analysis (WebGestalt), and their normalized gene expression signals are graphed along with their idealized templates (Figure 8).

**Table 4.**
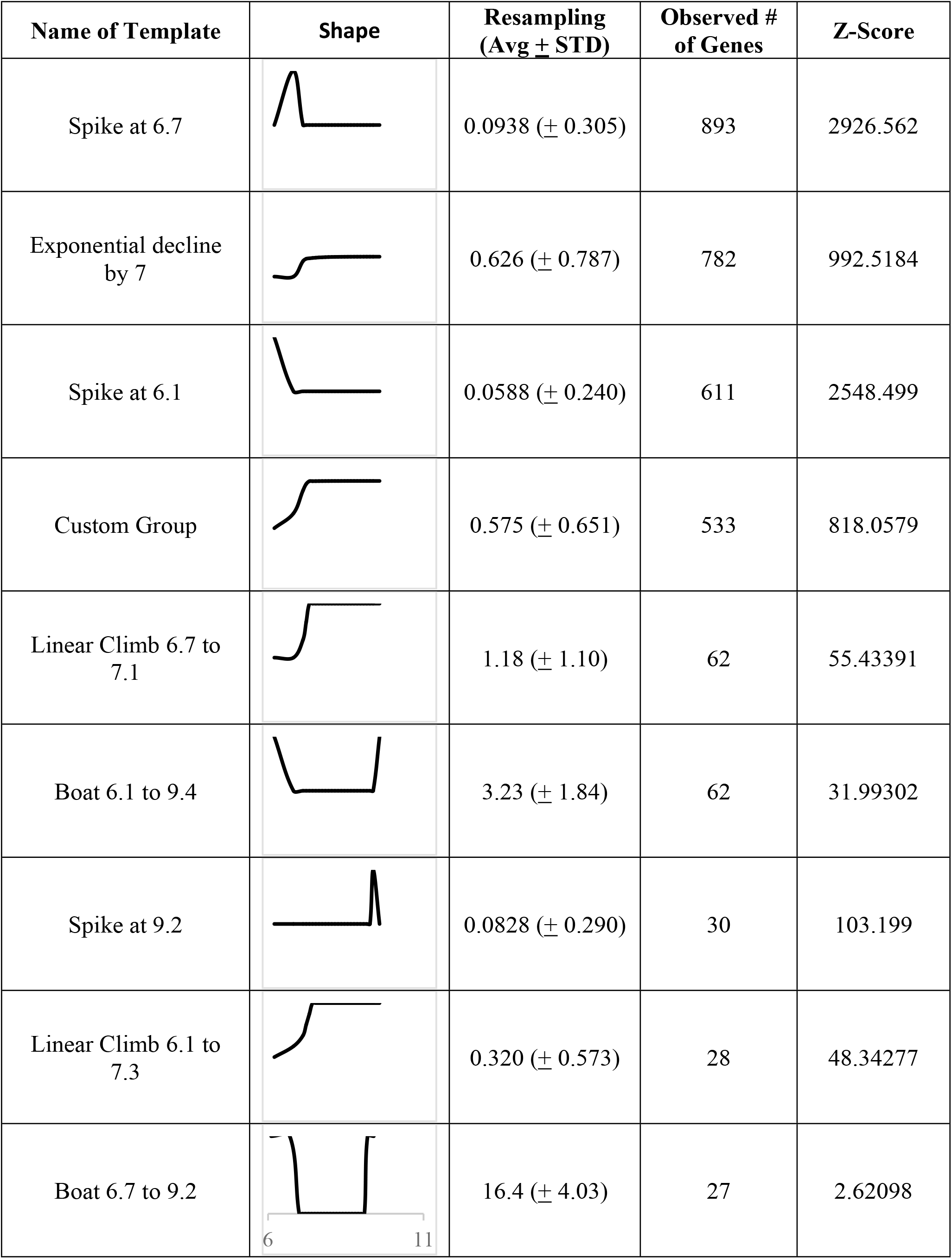
Templates to which gene expression most commonly fit. Each of these templates had more genes than expected by chance. The expected number of genes was found using a resampling (1000 iterations) and was considered to be significantly represented is Z-score ≥ 2.

| Name of Template | Shape | Resampling<br>(Avg ± STD) | Observed #<br>of Genes | Z-Score |
| --- | --- | --- | --- | --- |
| Spike at 6.7 | | 0.0938 (± 0.305) | 893 | 2926.562 |
| Exponential decline<br>by 7 | | 0.626 (± 0.787) | 782 | 992.5184 |
| Spike at 6.1 | | 0.0588 (± 0.240) | 611 | 2548.499 |
| Custom Group | | 0.575 (± 0.651) | 533 | 818.0579 |
| Linear Climb 6.7 to<br>7.1 | | 1.18 (± 1.10) | 62 | 55.43391 |
| Boat 6.1 to 9.4 | | 3.23 (± 1.84) | 62 | 31.99302 |
| Spike at 9.2 | | 0.0828 (± 0.290) | 30 | 103.199 |
| Linear Climb 6.1 to<br>7.3 | | 0.320 (± 0.573) | 28 | 48.34277 |
| Boat 6.7 to 9.2 | | 16.4 (± 4.03) | 27 | 2.62098 |

**Figure 8.**
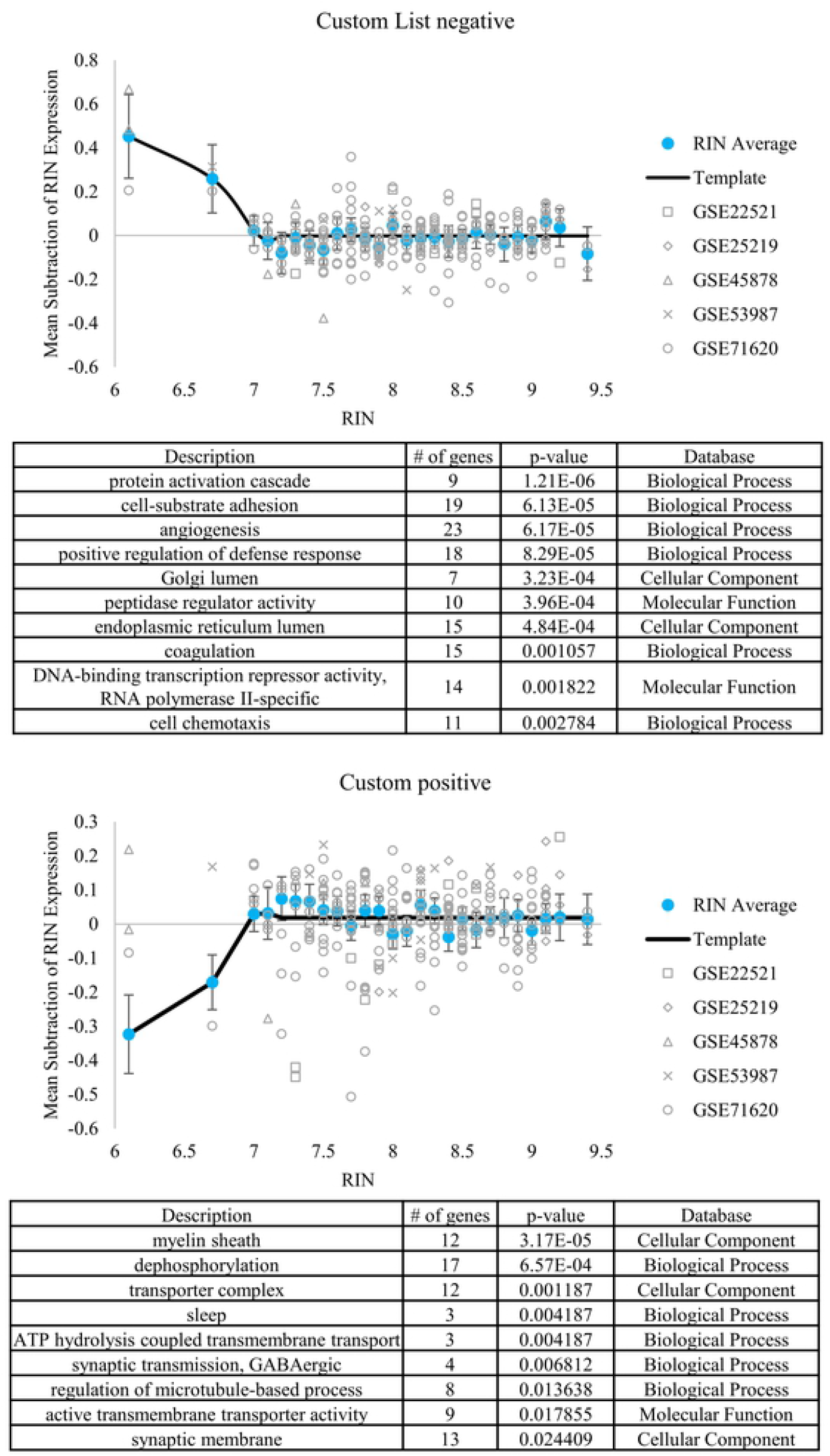
Representative gene expression templates. Each figure represents a template associated with a significant number of genes. The table under each figure is the overrepresented pathways corresponding to the template. Custom negatively correlated (A). Custom positively correlated (B). The blue dots represent average of gene expression. Each gray shape represents a subject, and each corresponds to a study. The black line represents the template.

The “completely linear” template in this phase of the analysis (the one in which gene expression steadily rises as RIN improves) was designed to replicate the relationship ostensibly found by the Pearson’s test in the independent data-subset (e.g., Figure 5), yet it failed to do so. This follows the earlier observation that the independent data-subset Pearson’s correlation, although it found a robust set of genes across multiple studies, did not appear to entirely capture the nature of the RIN-to-gene relationship. For instance, in the data-subset with the largest n (GSE71620, n = 199), the r-value range, while significant, was also restricted, and rarely rose above |0.34|. This suggests a pattern was detected by the Pearson approach, but it is not entirely linear. The overrepresented “exponential rise at 7” pathway reflected the monovalent inorganic cation transmembrane transport activity (GO: 0015077; 17 genes out of 420 genes; p = 5.05 × 10^−3^) found in the “independent data-subset” analysis, suggesting this may be a more accurate representation of a relatively narrow range over which RIN and gene expression are related. Therefore, we graphed the average gene expression for the genes present in the membrane-bound pathway as a function of RIN with the individual template superimposed (Figure 2D).

### Relationship between Alzheimer’s disease and RIN-sensitive genes

In order to compare RIN-sensitive genes to AD differentially expressed genes, two approaches were used. First, a comparison between two individual datasets, one reflecting RIN-sensitive and another reporting AD-associated genes was used. Second, consensus findings of RIN-sensitive vs. AD-associated genes from multiple datasets were compared. For the first approach, a common set of genes with sufficient signal intensity for testing was established between Miller et al (2017; (15)) and Chen et al (GSE71260; (37)). Among these 11,631 genes, Chen found 4,173 RIN-sensitive genes while Miller found 3,238 AD-significant genes. The log2 fold changes (L2FCs) for the ‘AD-effect’ and the r values for the ‘RIN-effect’ are plotted (Figure 9A) and revealed a strong directional agreement. More genes were found significant by both AD and RIN (1,531 genes significant by both AD and RIN; Data S4) than expected by chance (expect 1,162 genes, p < 1.0E-15), consistent with prior reports (15) noting the relationship between AD-associated and RIN-sensitive gene expression. Further, the direction of these changes appeared consistent, with genes downregulated with AD also showing a positive correlation with RIN (indicating gene expression decreases as RIN is lowered). Of the 1,531 genes significant by both RIN and AD, we determined if any directional groups (i.e. positive AD-associated genes and positive RIN-sensitive genes, positive AD-associated genes and negative RIN-sensitive genes) were overrepresented. In fact, genes whose expression was both decreased with AD and positively correlated with RIN, formed the majority of all commonly significant findings and were significantly overrepresented (n = 1223, binomial test, p < 1.0E-15; Figure 9A). This indicates the downregulated (although not upregulated) component of AD-associated gene expression is associated with RIN-based measures of RNA degradation.

**Figure 9.**
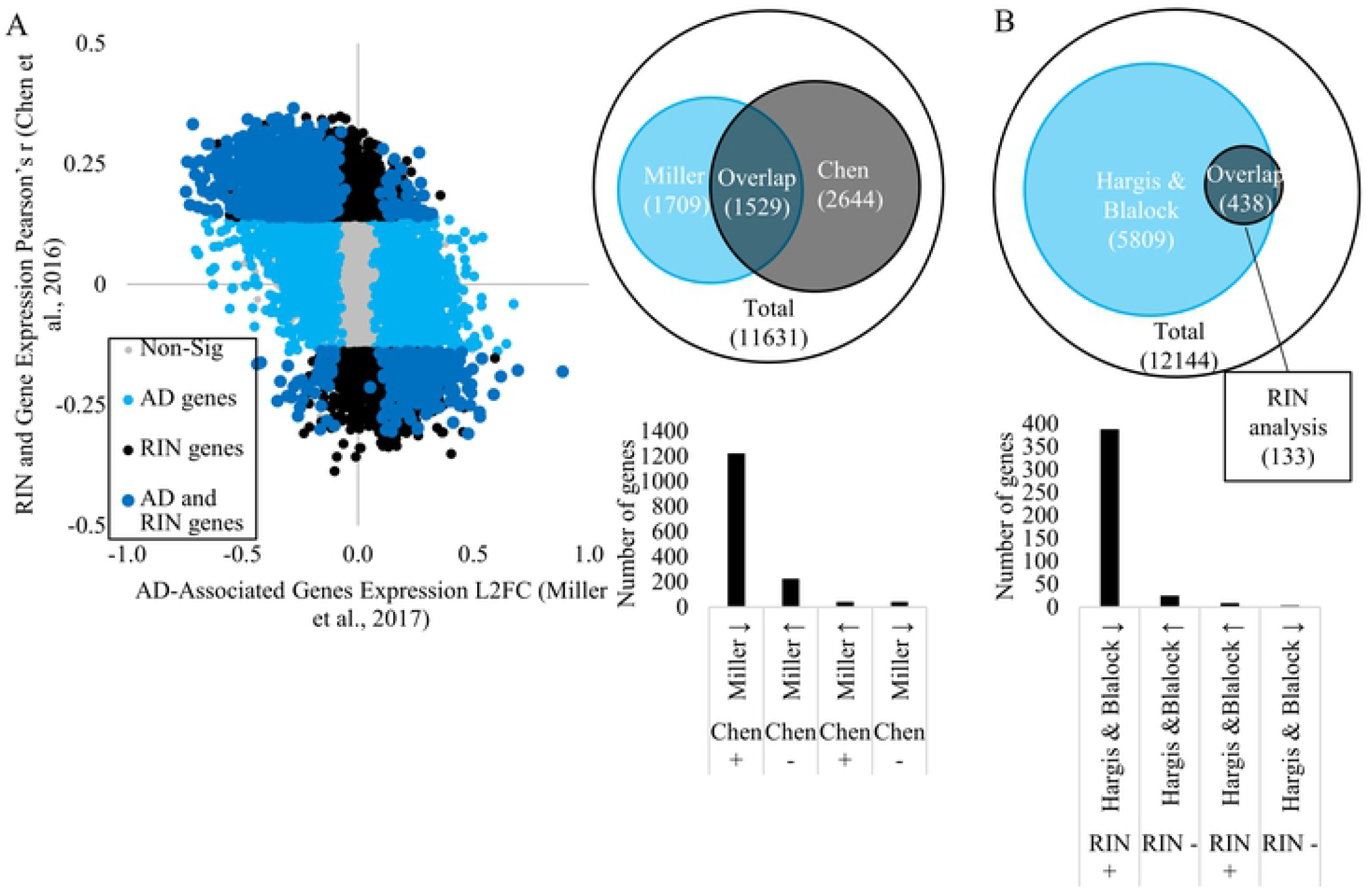
Relationship between RIN-sensitive genes and AD-associated genes. Single study comparison between the L2FC for AD genes (Miller et al., 2017) and correlation of RIN-sensitive genes (GSE71620; Chen et al., 2016). The Venn diagram also indicates the number of significant AD-associated genes, significant RIN-sensitive genes, and the overlap from the Miller et al. and Chen et al. analysis. The bar graph shows the number of significant overlapping genes and the Pearson’s correlation direction for RIN and direction of the L2FC (A). An analysis of the commonalities between a consensus of AD-associated genes (Hargis & Blalock, 2017) and the agreement of multiple RIN-sensitive datasets (Figure 5). The Venn diagram indicates the number of significant AD-associated genes, significant RIN-sensitive genes, and the overlapping genes. The bar graph shows the number of significant overlapping genes separated by the unanimous direction of the Pearson’s correlation for RIN and the average L2FC direction (B).

For the second approach, we then repeated this process, but used a more generalized approach by applying the Hargis and Blalock (2017; (1)) and current data (Figure 5) were indexed to find a common set of 12,144 genes. There were 6247 significant AD-associated genes and 571 RIN-sensitive genes (binomial p = 4.64E-08, p < 1E-15, respectively). There were more genes that were both AD-associated and RIN-sensitive than expected by chance as well (expected: 294; found: 438; binomial p= 6.66E-16; Data S5; Figure 9B). We found the majority of these genes (383/ 438) had expression that was both decreasing with AD and positively correlated with RIN (expected: 110; binomial p< 1.0E-15; Figure 9B), supporting a relationship between RIN-sensitivity and AD influence on gene expression, although the AD effect appeared to encompass more genes than could be explained by the RIN effect.

This analysis supports prior findings (15) that RIN is associated with gene expression levels, especially for AD-associated genes. However, our findings also support the hypothesis that the influence of RIN extends beyond AD and influences AD-associated gene expression even in normal, non-pathological brain tissue. This suggests AD may amplify some general property of RNA degradation present pre-mortem even in non-pathological samples.

## Discussion

RNA integrity numbers (RINs) have been important tools in determining RNA quality for nearly fifteen years, however the relationship between RIN and gene expression is still not well understood. Changes in RIN values may reflect the combined influence of multiple insults (e.g, RNAse activity, oxidative stress, tissue injury) (8, 48) to which different mRNA species may be selectively vulnerable. For the present work, we designed an analysis that tested individual, post-mortem, human frontal cortex data-subsets independently (secondary data analysis), and as a single dataset (meta-analysis). We hypothesized that, despite multiple possible mechanisms of insult, consistent genes and pathways would associate with RNA quality in post-mortem brain samples from control subjects with no diagnosed neuropathology.

Our findings support this by showing specific genes and pathways are consistently associated with RNA degradation across multiple studies. However, while we hypothesized a linear relationship between RNA integrity numbers (RINs) and gene expression, our template analysis found that an exponential (or possibly sigmoidal) relationship better describes individual gene relationships to RIN. In addition, we found a surprising difference between RNA degradation’s influence with sex and age (aged males tended to show the strongest relationship). Based on the present work and prior publications (15), we propose that the apparently neuron-selective RNA degradation in control tissue is the result of selective damage to mRNA localized in synapses, and is similar to the influence seen in Alzheimer’s disease (AD).

Regarding other metavariables commonly used to assess tissue quality, RIN values in healthy control brain tissue appears to correlate inconsistently with post-mortem interval (PMI) or tissue pH. Since our analysis does rely on the information published by others, there could be selection criteria we are unaware of that limit the range of variables. This ‘floor-effect’ could be impacting our results. With the information available to us, only 2 out of the 5 datasets report PMIs with a significant correlation between PMI and RINs, and they correlate in opposite directions. This is particularly of note since several studies investigating the influence of RIN in controlled experimental settings have manipulated PMI to drive changes in RIN (9, 49), which may represent a component of the process observed here or an entirely separate process. Pre-mortem changes in control tissue that are not associated with known pathology or mishandling may also play a role. A study from the University of Maryland used human brain cortices from the NIH found the quality of RNA did not significantly drop until 36 hours post-mortem (50). With the exception of subjects from GSE46706, the longest PMI for any subject was 37.5 hours; this may explain why there is not a clear relationship between PMI and RIN in the present work. Our analysis is in line with others (51-53) that have found a lack of a relationship between RIN and PMI (however, see (54)). It has also been established the PMI and RIN correlation is tissue-dependent, and possibly region-specific as well (3). One study found subjects with Alzheimer’s disease had a significant negative correlation between PMI and RIN, but the control subjects did not (55). Therefore, more investigation needs to be done on the relationship between PMI and RNA quality. Like PMI, tissue pH does not show a robust relationship with RIN in control brain tissue in this study. Only one out of the four studies reporting pH finds any significant correlation. However, other studies have found a significant relationship between RIN and pH (18, 56, 57) and it is possible, since pH is also used as a preliminary triage variable, that the work we analyzed had a constrained pH range. It should be noted that even though these metavariables did not have a robust impact on RIN, on their own they did show a significant association with transcriptional profile (albeit different sets of genes). This suggests these metavariables may have mechanistically distinct influences. Because of this, PMI and pH appear to be unreliable proxies for RIN-assessments.

In females, younger subjects are moderately more sensitive to RNA degradation than aged. But in males, there is a strong increase in RIN-sensitivity with age (Figure 4B). Age itself does not significantly correlate with RIN, even when young males, young females, aged males, and aged females from the GSE71620 data-subset are analyzed separately. One possibility is that aging genes could be indirectly influencing RNA degradation since aging can play a role in mRNA turnover (58). Future work could focus on whether aging itself confers a differential sensitivity of mRNA to RIN variability, and what the potential mechanisms of such an effect would be and what the potential mechanisms of such an effect would be.

Our data indicates RNA degradation is strongly associated with specific genes and pathways in post-mortem human frontal lobe tissue in control subjects. We find the same outcomes using two types of analyses: comparing individual datasets and merging datasets for a single analysis. In both cases, there is a highly consistent subset of genes that reliably showed lower expression as a function of decreasing RIN. The functional pathways with which these genes are associated included vesicles, transport activity, synapses, and mitochondria, implying neuronal mRNA may be particularly sensitive to degradation.

There are two thresholds to consider for RIN’s influence on mRNA expression-1) where RIN begins to have a significant effect on gene expression; and 2) where RIN becomes so severe that gene expression reaches a floor effect and is no longer impacted by further RNA degradation. Although individual RIN-sensitive gene sensitivities vary for both of these thresholds, there are some general consensus values that may be useful to consider in future work. As RNA degrades and the RIN begins to decrease, there is a marked increase in the number of genes whose expression correlates with RIN (Figure 6). This indicates that if experimental tissue has a RIN above 8.6, then in general, there is no reliably detectable effect of RIN on gene expression and RIN correction would not be necessary in human post-mortem neocortical tissue. Between RINs of 7.2 and 8.6, there is a strong increase in RIN-sensitive mRNA detection, and below 7.2, detection again falls below statistically significant thresholds. However, our analysis also suggests that specific pathways (e.g., monovalent cation transporter-Figure 2) are more impacted at lower RINs. Thus, it is possible that different pathways are impacted at different points along the continuum of mRNA degradation.

In this work, RNA degradation also appears to exert a floor effect on gene expression. The point at which a gene reaches its floor effect can vary from gene to gene, but is critical to consider. Beyond this threshold, the application of a RIN correction procedure would appear to provide data but instead would provide nonsense as its starting value cannot be estimates based on the RIN score. Often, a RIN of 6 is used as a criterion for inclusion in a study, though there is no clear consensus in the literature (56). Based on our findings using 6 data-subsets, a RIN of 6.7-7 appears to represent a generalized ‘point of no return’ for RIN-sensitive genes. This floor effect is also indicated by GSE46706, a data-subset with an unusually high RIN-to-gene expression false discovery rate (FDR) > 0.75 and an unusually low range of RIN values (2-7.7; only 26 of the 119 subjects had a RIN ≥ 6). This supports our hypothesis that lower RINs are associated with flattened gene expression, and suggests that low range RIN corrections are unreliable for expression adjustment.

Taken together, the upper and lower thresholds for RIN-sensitivity suggest that there is an interior RIN range between 7.2-8.6 in which RIN may have a fairly linear effect on gene expression that is amenable to correction. However, when considering RIN-sensitive genes, for RIN values above this range, RIN correction may inappropriately add to variance, since these genes do not need correcting. For RIN values below this range, we argue that the signal is not rescuable.

Although there is strength to secondary and meta-analyses such as the present work, especially regarding robust findings, there are important caveats. These findings are correlative, and therefore we cannot determine if the associations are causal, consequential, or epiphenomenal. Further, while it is tempting to speculate that RNA degradation is impacting genes/ pathways at different rates (perhaps even forming a ‘meta-pathway’ of progressive decay), this would require more observation and interventional experimentation. In addition, there is evidence that different pathways regarding mRNA transcription and transport are impacted at differing rates in post-mortem mice and zebrafish (4), giving more credence to the possibility that RNA degradation results in a cascading effect on the transcriptome. Although it is expected that available datasets would have restricted RIN ranges (RIN is typically used to establish a quality control cutoff; similar issues exist for other metavariables like pH and PMI), a more complete idea of RNA degradation’s effects could be achieved by examining the transcriptional influence across the entirety of the RIN range and with more attention given to the different measures (e.g. Total RNA ratio, 28S peak height, fast area ratio, and the marker height) that contribute to the RIN score (11). However, another consideration is the subject-to-subject normal variation, and RIN may not reach such low values in the absence of severe pathology.

Our findings that pathways relating to neurons are most vulnerable to RNA degradation suggests concerns with use of common regression approaches to control for RIN in transcriptional profiling of neural tissue, particularly with regard to neurodegenerative disease. First, we report a consistent set of genes sensitive to both RIN and Alzheimer’s disease (AD). These RIN/AD genes predominantly have lower expression with worsening AD and with lower RNA quality (e.g., are decreased with AD and show a positive correlation with RIN). These results support Miller et al., where RIN correction procedures removed the apparent Alzheimer’s disease effect (15). Miller et al. reported RNA degradation was a confounding variable worsened in AD samples, and proposed this might be due to improper handling of tissue (15). However, questions remain regarding whether it is reasonable to assume that pathologists would be selectively indelicate with AD compared to control tissue, and it should be considered that the RNA could decay appreciably in living tissue prior to death and independent of post-mortem procedures.

Second, in a disease state where both RIN and AD gene expression are declining, attempting to control for one issue may inadvertently remove clinically relevant findings from the other. We found similar results when we compared an AD dataset consisting of the robust AD changes observed in multiple independent studies (1) along with the consistent RIN-sensitive genes identified in 3 control tissue data-subsets (see cross-dataset analysis using individual datasets in Results). This supports the observation (15) that specific AD genes are RIN sensitive. This may indicate correcting and/or normalizing RIN may actually be removing a true effect within AD subjects, as the RIN degradation itself may reflect in situ degradation, not tissue mishandling.

There are still many unanswered questions regarding the RNA degradation and its impact on gene expression. To our knowledge, this is the first study to look across multiple datasets at the relationship between RIN and gene expression in human post-mortem control brain tissue. However, other studies have looked at RNA degradation’s impact in other tissues and species. For example, when Opitz et al. investigated RNA degradation in RNA extracted from renal cancer tumors, they found overrepresented translation, GTP, and RNA activity (7) pathways. Gallego-Romero et al. found inflammatory, immune, and clotting factors are most vulnerable to rapid RNA degradation in human peripheral blood mononuclear cells (10). Jaffe et al. also presented evidence that RNA from different cell types may have different susceptibility to degradation (9). This indicates cell and tissue type plays a role in which pathways are sensitive to RNA degradation. Indeed, this suggests that the mRNA associated with the principal functional cells of a tissue are the most vulnerable to degradation.

In conclusion, transcriptional profiling reveals that neuronal, especially synaptic, pathways appear more strongly and consistently associated with RNA integrity than other pathways in control, post-mortem, human frontal lobe tissue from multiple independent datasets. This suggests that mRNA localized to synaptic regions may be particularly vulnerable, possibly due to its close association with mitochondria. Further, RNA integrity number (RIN) does not reliably correlate with two other measures commonly associated with post-mortem tissue quality, PMI and pH, suggesting PMI and pH would be poor proxies for RIN. Interestingly, there does appear to be an age-based shift in RIN-sensitive gene expression.

Despite the fact that the range of RIN values was roughly the same in young and aged subjects, the association between RIN and gene expression increased markedly with age in males, and to a much lesser extent, decreased with age in females. In addition, and in support of prior work (15), there is strong overlap between RIN and AD’s influence on gene expression. For example, we also found that nearly half of all AD-sensitive genes identified in Miller et al. (15) were also correlated with RIN (in Chen et al. (37), completely independent dataset of control-only post-mortem brain tissue). In datasets where RIN and AD are confounded (e.g., where RIN is significantly lower in AD samples), procedures such as multiple regression that are designed to correct for RIN’s influence may inadvertently remove the effect of AD. Indeed, one of AD’s effects may be to lower RIN. Therefore, we suggest avoiding RIN-correction if such a confound exists, as have prior researchers (10). Reporting the confound may be more appropriate than obfuscating it with RIN normalizing procedures. We also found that the relationship between RIN and gene expression more closely reflects a sigmoidal (or stepped) relationship across a narrow RIN range of 6.7-8.6, rather than a continuous linear relationship across the entire RIN range. This suggests that RIN correction tools using regression may over-correct values outside of the linear range, and under-correct values within it.

